## Supplemental Material for "Screening of metabolism-disrupting chemicals on pancreatic α-cells using in vitro methods"

**Supplementary Table 1. List of antibodies used in this study.**

| **Target antigen** | **Antibody Name** | **Manufacturer and catalogue number (Cat no.)** | **Species raised in** | **Dilution** | **RRID** |
|---|---|---|---|---|---|
| BiP | BiP Antibody | Cell Signaling Technology; Cat no. 3183 | Rabbit, polyclonal | 1:1000 | AB_668355 |
| p-eIF2α | Phospho-eIF2α (Ser51) (119A11) | Cell Signaling Technology; Cat no. 3597 | Rabbit, monoclonal | 1:1000 | AB_390740 |
| α-Tubulin | Monoclonal Anti-αTubulin antibody | Sigma; Cat no. T9026 | Mouse, monoclonal | 1:5000 | AB_477593 |
| Goat anti-mouse IgG | Goat Anti-Mouse IgG (H+L) HRP Conjugate antibody | Bio-rad; Cat no. 170-6516 | Goat, Polyclonal | 1:5000 | AB_11125547 |
| Goat anti-rabbit IgG | Goat Anti-Rabbit IgG (H+L) HRP Conjugate antibody | Bio-rad; Cat no. 170-6515 | Goat, Polyclonal | 1:5000 | AB_11125142 |

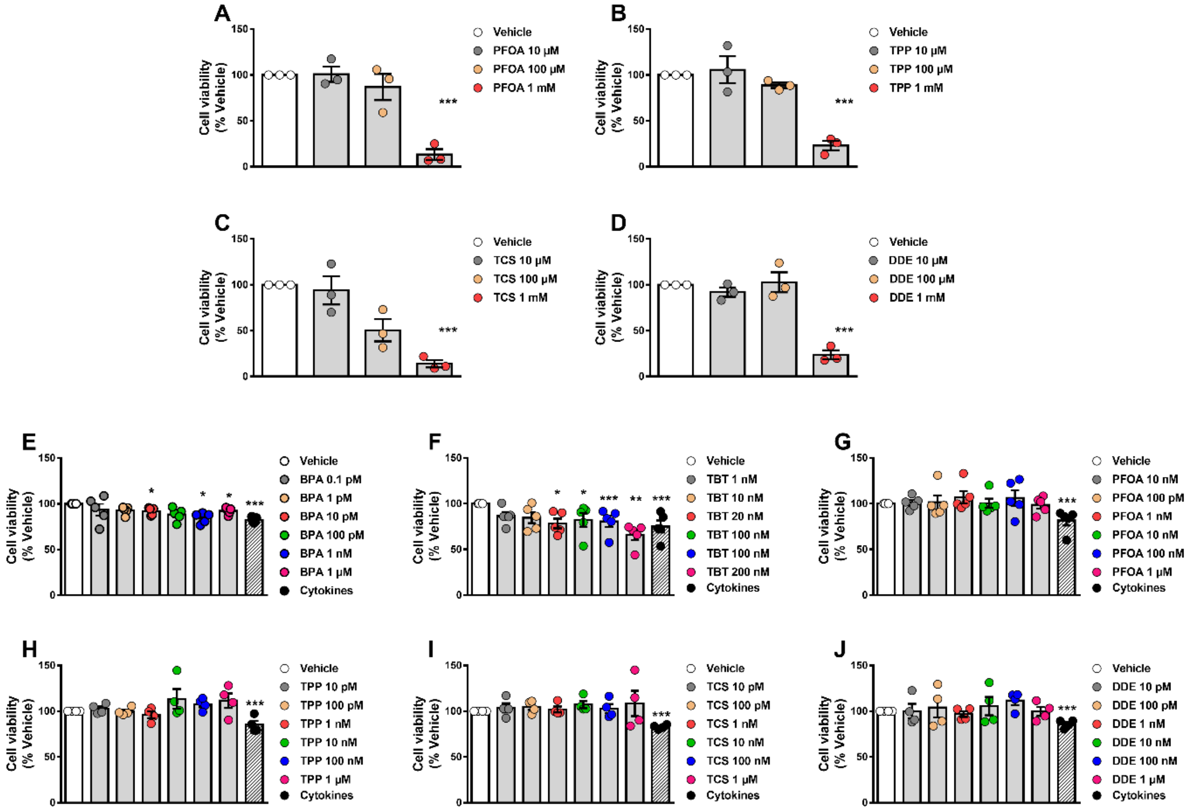


**Supplementary Figure 1. α-cell viability upon MDC exposure.** (A-D) αTC1-9 cells were treated with vehicle (DMSO) or different doses of PFOA (A), TPP (B), TCS (C), or DDE (D) for 48 h. (E-J) αTC1-9 cells were treated with vehicle (DMSO) or different doses of BPA (E), TBT (F), PFOA (G), TPP (H), TCS (I), or DDE (J) for 72 h. A cocktail of the cytokines IL-1β + IFNγ (50 and 1000 U/ml, respectively) was used as a positive control. Cell viability was evaluated by MTT assay. Results are expressed as % vehicle-treated cells. Data are shown as means ± SEM (n = 3-5 independent experiments, where each dot represents an independent experiment). * $p \leq 0.05$, ** $p \leq 0.01$ and *** $p \leq 0.001$ vs. Vehicle. MDCs vs. Vehicle by one-way ANOVA; Cytokines vs. Vehicle by two-tailed Student's *t* test.
